# *Cis*-regulatory code for predicting plant cell-type specific high salinity response

**DOI:** 10.1101/466326

**Authors:** Sahra Uygun, Christina B. Azodi, Shin-Han Shiu

**Affiliations:** Genetics Program, Michigan State University, East Lansing, MI, USA; Department of Plant Biology, Michigan State University, East Lansing, MI, USA; Department of Computational, Mathematics, Science, and Engineering, Michigan State University, East Lansing, MI, USA

**Author notes:** Corresponding author: Shin-Han Shiu, Michigan State University, Plant Biology Laboratories, 612 Wilson Road, Room 166 East Lansing, MI 48824-1312. Sahra Uygun Agendia Inc. Irvine, CA 92618, USA. These authors contributed equally to this work (Sahra Uygun, Christina B. Azodi).

## Abstract

Multicellular organisms have diverse cell types with distinct roles in development and responses to the environment. At the transcriptional level, the differences in environmental response between cell types are due to differences in regulatory programs. In plants, although cell-type environmental responses have been examined, details on how these responses are regulated remain spotty. Here, we identify a set of putative *cis*-regulatory elements (pCREs) enriched in the promoters of genes responsive to high salinity stress in six *Arabidopsis thaliana* root cell types. Using machine learning with pCREs as predictors, we establish *cis*-regulatory codes, *i.e.* models predicting whether a gene is responsive to high salinity for each cell type. These pCRE-based models outperform models utilizing in *vitro* binding data of 758 *A. thaliana* transcription factors. Surprisingly, organ pCREs identified based on whole root high salinity response can predict cell-type responses as well as pCREs derived from cell-type data -because organ and cell-type pCREs predict complementary subsets of high salinity response genes. Our findings not only advance our understanding of the regulatory mechanisms of plant spatial transcriptional response through *cis*-regulatory codes, but also suggest broad applicability of the approach to any species, particularly those with little or no *trans* regulatory data.

## Introduction

The identification of different types of cells and the characteristics that make them unique in multicellular organisms has fascinated and challenged biologists since Anton van Leeuwenhoek’s invention of microscope in the late 17^th^ century (1). These distinct cell types carry out, to various degrees, specialized functions that contribute greatly to organismal complexity. One of the crucial components that allows for these specialized functions is differences in transcription regulatory mechanisms, which allow for cell-type specific gene expression during development as well as in response to changing environmental conditions. To study cell-type specific gene expression profiles, isolation of individual cell types is required because the gene expression levels might not reflect per cell-type changes if a whole organ is analyzed (2). Two prominent approaches for isolating distinct cell types include fluorescent activated cell sorting and laser capture microdissection, both of which have been applied to multiple metazoan species including *C. elegans*, Drosophila, mouse and human (3–8), as well as plants (9–12). In plants, root is an ideal system to study cell types as the roots have radial organization with layers of distinct cell types and undergo continuous development since their derivation from stem cells (13). In addition, the cell-sorting-based approaches have been developed to study cell-type specific expression in *A. thaliana* root development (10, 14) and nitrogen/high salinity responses among root cell types (9, 15, 16). These studies of root cell types significantly advance our understanding of how individual cell types differ in gene expression over time and in response to different environmental conditions, including high soil salinity, which results in reduced yield in crops (17).

While there is an understanding of how root cell types differ in their transcriptional response to high salinity (9), it remains a major question how such cell-type specific response is regulated via *cis*-regulatory elements (CREs), transcription factors (TFs), cofactors, and chromatin remodeling complexes (18). At the *cis*-regulatory level, multiple studies have utilized cell-type specific data to globally identify CREs underlying differential gene expression across cell types in metazoans (19–23). A similar study in plants is now feasible for two reasons. First is the availability of *A. thaliana* global *in vitro* TF binding data generated with Protein Binding Array (PBM) and DNA Affinity Purification (DAP) (24, 25). Second is the availability of computational methods for identifying putative *cis*-regulatory elements (pCREs) (26–30), which have facilitated the identification of stress related pCREs and those contributing to organ-specific stress response (31, 32). These CREs can be used further to establish a stress *cis*-regulatory code with machine learning (31), i.e. a computational model that answers the question how and to what extent a set of CREs collectively control transcriptional response under a stress condition. Our recent studies on *A. thaliana* provided spatial *cis*-regulatory codes of stress responsive gene expression at the organ level (root vs. shoot) (32). Currently, there is no *cis*-regulatory code available that explain stress response at individual cell-type level.

The regulatory mechanisms responsible for plant cell-type specific responses to external factors remain largely unknown (33). In this study, we aimed to investigate the *cis*-regulatory code of high salinity responsive gene expression (particularly up-regulation) in six root cell types using an existing dataset (9). First, we asked to what extent high salinity responsive gene expression is similar between whole root and the individual root cell types. Next, we assessed the extent to which large-scale *in*-*vitro* TF binding information (24, 25) as well as organ-specific pCREs (32) could predict root cell-type high salinity up-regulation. Furthermore, we identified pCREs that likely regulate high salinity up-regulation in each cell-type and established cell-type *cis*-regulatory codes.

## Methods

### Gene expression datasets and their processing

The root cell-type high salinity stress expression dataset (9) was downloaded from Gene Expression Omnibus (GSE7641). This expression dataset consists of control and high salinity stress conditions (150mM NaCl treatment for 1h) for the following cell types: columella (COL), cortex (COR), endodermis/quiescent center (END), epidermis (EPI), proto-phloem (PHL) and stele (STE). The Affymetrix CEL files were pre-processed and quantile normalized using the Bioconductor affy package in R environment (https://bioconductor.org/packages/release/bioc/html/affy.html). For differential gene expression, log_2_ fold changes and associated *p*-values were calculated using high salinity stress treatment and corresponding control samples for each cell-type with the limma package from Bioconductor (34). The *p*-values were adjusted for multiple testing (35). The whole root abiotic stress dataset from AtGenExpress (http://www.weigelworld.org/resources/microarray/AtGenExpress/) was also used and processed according to a previous study (31). A gene was considered up-regulated in a cell-type or in the whole root if its log_2_ fold-change value was ≥ 1 and the adjusted *p*-value was ≤ 0.05. Non-responsive genes were defined as genes that were neither up-nor down-regulated under any stress at any time point in any sample from the AtGenExpress data as defined previously (32).

### Gene Ontology (GO) enrichment analyses

To find functional categories that were significantly over- or under-represented in the organ and root cell-type up-regulated genes, GO slim terms were retrieved (http://www.geneontology.org/ontology/subsets/goslim_plant.obo). The genes annotated to each GO term that were high salinity up-regulated (in root, shoot or one of the six root cell types) were compared against the rest of genes in the same GO term to build a 2×2 contingency table. Enrichment was tested with the Fisher’s exact test. The *p*-values of enrichment were adjusted for multiple testing with the *q*-value method (36). The enrichment score was reported as −log(*q*-value). The same approach was used to identify functional differences between STE genes (A) correctly or incorrectly predicted by both cell-type and all organ pCREs and (B) correctly predicted by only cell-type or organ pCREs.

### Gene co-expression analyses

To find co-expressed gene clusters, the root cell-type high salinity stress expression dataset was combined with the root stress expression dataset from AtGenExpress (37). Genes in the combined dataset were classified into co-expression clusters using c-means (38) in the R environment. Among the resulting clusters (**File S1**), those with < 10 genes were excluded from further motif finding analyses because the motifs identified would have limited statistical support. The clusters with > 60 genes were further divided using c-means, resulting in 538 clusters included for further analysis, each with 10-60 genes (**File S1**). This range of number of genes in a cluster was required for efficiently running the motif finders (31). Fisher’s exact test was used to identify the clusters with over-represented numbers of high salinity up-regulated genes in each root cell-type compared to the rest of the genome (multiple testing q ≤ 0.05) (36).

To determine if STE high salinity up-regulated genes that were predicted correctly by different sets of pCREs had different expression patterns, STE genes were clustered into 8 clusters using *k*-means (fcp R package: https://cran.r-project.org/web/packages/fpc/fpc.pdf) in the R environment. Next, genes that were correctly predicted by both, either, or neither cell-type and organ pCREs were tested for enrichment of genes from each of the 8 clusters using the Fisher’s exact test and multiple testing correction described above. The same approach was used to identify differences in the presence or absence of organ and cell-type pCREs between STE genes that were predicted correctly by different sets of pCREs. Only the top 100 most important pCREs from each set, as determined by our machine learning models, were used for clustering.

### Analysis of TF binding data

To identify whether existing *in*-*vitro* TF binding data could explain root cell-type high salinity responsive gene expression, two sets of TF binding datasets were obtained. These datasets included Position Frequency Matrices (PFMs) obtained from the CIS-BP database (24) and DAP-seq peaks (~200 bp long) obtained from the *A. thaliana* Cistrome study (25). The handling of the TF binding data was as described previously (32). Both CIS-BP and DAP-seq information were used in predicting cell-type high salinity up-regulation. CIS-BP PFMs were converted to position weight matrices (PWMs) and mapped to *A. thaliana* promoter sequences. Similarly, DAP-seq peak sites that correspond to *A. thaliana* promoters were used in predictions. Note that the promoters were defined as the regions 1,000bp upstream of transcription start sites.

### Identification of pCREs regulating cell-type high salinity response

To identify pCREs relevant to high salinity up-regulation in a cell-type from the promoter regions, a previously established pipeline (31) was applied to each co-expression cluster enriched in high salinity up-regulated genes in a given cell-type. This pipeline tested if a pCRE, *X*, was significantly more likely to be found in the promoter regions of genes up-regulated in cell-type *C*, compared to the promoters of non-responsive genes, using Fisher’s exact test. To evaluate the impact of threshold significance levels in calling a pCRE as over-represented, we applied two threshold *q*-values at 0.05 and 10^−6^ that lead to 7,417 and 3,095 pCREs (**File S2, S3**) enriched in at least one cell-type, respectively. As the prediction performances (see Predictive models of cell-type high salinity up-regulation) were similar using 7,417 (AUC-ROC = 0.71-0.79) and 3,095 (AUC-ROC = 0.68-0.76; **Fig. 3A, Table S1**) pCREs. The pCRE set with a more stringent enrichment threshold was used in further analyses. To assess similarity between pCREs, a Pearson’s Correlation Coefficient (PCC) was calculated using the Position Weight Matrices (PWMs) of a pCRE pair as described previously (31).

### Predictive models of cell-type high salinity up-regulation

To establish the *cis*-regulatory code for genes up-regulated by high salinity treatment in each cell-type, we built a machine learning model capable of predicting whether a gene would be up-regulated or non-responsive for each of the cell types of interest. To assess the impact of machine learning methods in building such models, Support Vector Machine (SVM, (39)) and Random Forest (RF, (40)) were tested using the Waikato Environment for Knowledge Analysis (WEKA, (41)).

To find the optimal parameters for classification, grid-searches were performed. The parameters for SVM were: 1. the ratio of non-responsive to up-regulated genes, 2) the soft margin, and 3. the gamma parameter of the Radial Basis Function kernel. The RF parameters included: a. the ratio of non-responsive to up-regulated genes, and b. number of features (i.e. pCREs) to use in trees. A standard 10-fold cross validation scheme was used to prevent model overfitting. Two measures were used to evaluate the prediction performance. The first was the Area Under Curve-Receiver Operating Characteristic (AUC-ROC) measure, where a perfect model would have AUC-ROC = 1 and random predictions would lead to AUC-ROC = 0.5. The second approach was plotting the precision-recall curve, where precision was the ratio of true positive predictions to overall number of genes that were predicted as positive and recall was the ratio of true positive predictions to total number of positive class (high salinity up-regulated genes in a cell-type). The models with satisfactory classification would have precision-recall curves towards the upper-right corner of the graph and the models with random predictions would be no better than the background of ratio of positive to negative class. To assign importance scores to the features, SVM models were built using Scikit-Learn (42) and the absolute value of the coefficients assigned to each feature over 100 replicates were averaged. A gene was considered correctly classified by the model if its median predicted probability score over the 100 replicates was greater than the decision threshold.

## Results

### Comparison of organ and cell-type transcriptional response to high salinity

In *A. thaliana*, differential gene expression in organs and cell types has been studied in a genome-wide manner across developmental stages as well as in response to a variety of environmental stresses (9, 14, 37). Here we used an *A. thaliana* cell-type transcriptome data under high salinity stress (9) to dissect the *cis*-regulatory code driving cell-type response to stress. The six root cell types included columella (COL), cortex (COR), stele (STE), protophloem (PHL), epidermis (EPI) and endodermis (END). First, we asked to what extent the root cell-type high salinity responses differed from the whole organ high salinity response (32). We used whole organ (shoot or root) gene expression data focusing on abiotic and biotic stress treatments over multiple time points (37). To compare the global gene expression between samples, we calculated the between-sample Pearson Correlation Coefficient (PCC, **Fig. 1A, File S4**).

**Fig. 1.**
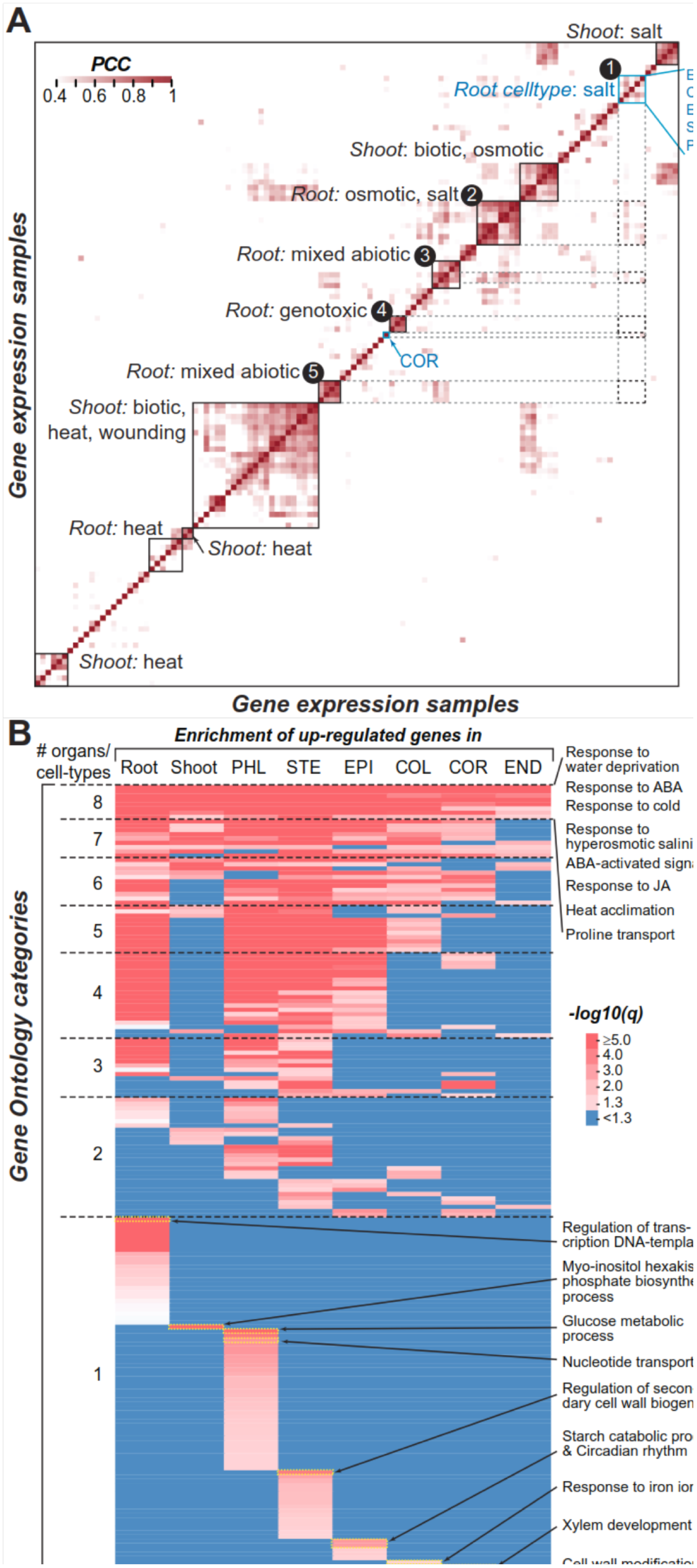
Gene expression correlation across stress datasets of root, shoot and root cell types. A. Heatmap of Pearson’s Correlation Coefficient (PCC) calculated using expression values of all sample pairs. The colors represent PCC values from low (lighter red) to high (darker red). PCC values < 95^th^ percentile of all pair-wise sample PCCs (0.42) are in white. Boxes with black outline are the clusters of similar treatments (e.g. root high salinity treatment samples clustering with root osmotic treatment samples). Boxes with blue outline and blue text indicates root cell-type samples including columella (COL), cortex (COR), stele (STE), proto-phloem (PHL), epidermis (EPI) and endodermis (END). Boxes with dashed black outline and gray dotted lines are for emphasizing the relationships between root cell-type high salinity treatment samples with whole root abiotic stress treatment samples.
B. Gene Ontology (GO) categories with overrepresented numbers of high salinity up-regulated genes in root, shoot, and/or root cell types. Dotted lines separate categories enriched in different numbers of organs/cell types. Shades of red: significant overrepresentation with *q*-value ≤ 0.05. Blue: *q* -value > 0.05. ABA: abscisic acid. JA: jasmonic acid.

The overall gene expression patterns of genes in high salinity-treated root cell types (box 1, **Fig. 1A**), except cortex (COR), were more similar to each other than to those of whole root and shoot (**Fig. 1A**), consistent with earlier findings (9). Additionally, a subset of the high salinity-treated root cell types were significantly and positively correlated with the high salinity/osmotic stress-treated whole root treatments (samples in box 2 and 3, **Fig. 1A**). This was not the case for other treatments (box 4 and 5, **Fig. 1A**). Given the cell types examined are a subset of cells examined in the whole root samples, the observed correlation between root cell-type and whole root responses is expected. However, we should emphasize that, confirming earlier studies (9), the degrees of correlation also indicated extensive differences in high salinity responsive gene expression between different cell types (e.g. PC_CCOR-PHL_=−0.05, PCC_EPI-PHL_=0.26) and between root cell-type and whole root samples treated for 0.5 or 1 hour (e.g. PCC_COR-ROOT_0.5h_ = 0.17, PCC_COR-ROOT_1h_=0.22). Thus, there is clearly information captured in cell-type data that cannot be obtained if only whole organ data are considered.

### Functional enrichment among spatially specific high salinity up8regulated genes

To explore the differences between high salinity-treated cell-type and whole root expression data further, we asked what Gene Ontology (GO) terms tend to be found among the high salinity up-regulated genes in root cell types compared to whole root and shoot up-regulated genes (**Table S2**). Eight GO terms were commonly over-represented among up-regulated genes in all organs and cell types (**Fig. 1B, Table S2**), including those relevant to abscisic acid-activated signaling, response to water deprivation and hyperosmotic salinity, proline transport as well as response to other stresses including cold and heat. These results suggest that, despite the substantial differences in their transcriptional programs (**Fig. 1A**), similar biological processes relevant to high salinity stress responses are activated regardless of the organ or cell-type. That said, there were also substantial differences in enriched GO terms specific to a subset of or to each organ/cell-type response. Among the 187 GO terms significantly overrepresented, 46% of them were specific to one gene set (**Fig. 1B, Table S2**). The finding that different biological processes are over-represented in root cell-type up-regulated genes is consistent with earlier studies indicating that spatial transcriptional response to stress is tissue-specific (9).

For example, glucose catabolic process was only enriched among phloem (PHL) high salinity up-regulated genes (**Fig. 1B**), reflecting the importance of phloem in transporting photosynthetic products (43) and the need to alter glucose metabolic gene expression in response to high salinity stress. In the columella (COL) genes, more iron responsive genes were up-regulated under high salinity stress (**Fig. 1B**), likely due to the connection between high salinity soil and reduced bioavailability of iron (44). Consistent with high salinity induced secondary cell wall thickening (45) and the proposed function of suberin in high salinity tolerance (46), genes relevant to the regulation of secondary cell wall biogenesis and cell modification genes involved in abscission (deposition of suberin/lignin) were up-regulated specifically in the stele (STE) and the endodermis (END), respectively (**Fig. 1B**). Another example was the starch metabolism genes specifically enriched among epidermal and lateral root cap (EPI) high salinity up-regulated genes (**Fig. 1B**). These starch granules could potentially bind to and reduce root sodium ion levels as reported in other plant species (47). Interestingly, circadian rhythm genes tend to be up-regulated specifically in epidermal cells as well; suggesting root circadian response may predominantly take place in the epidermis. It is not entirely clear why xylem development genes were enriched in high salinity responsive genes in the cortex (COR, **Fig. 1B**). The cortex cells may develop thickened secondary wall to counter high salinity stress. However, we cannot rule out the possibility that the procedure for isolating individual cell types introduce altered expression patterns, among other possibilities.

### Predicting cel8type high salinity up8regulation with Iarge8scale in8vitro TF binding data

Given the regulatory program responsible for controlling cell-type response to stress, not just under high salinity, remains largely unknown, we first focused on identifying TFs likely regulating high salinity up-regulation in each root cell-type. Extensive binding data for 758 *A. thaliana* TFs are available from two large-scale *in*-*vitro* TF binding studies, CIS-BP (24) and DAP-seq (25). We first tested which TFs might control cell-type gene expression under high salinity stress by identifying TFs with over-represented numbers of DAP-seq binding sites in the promoters of high salinity up-regulated genes in each root cell-type. Among cell types, 0-140 binding sites of TFs were enriched (Fisher’s exact test, *q* ≤ 0.05; **Table S3**). For COR and PHL, no sites were significantly enriched. On the other hand, COL and END cell types had the most TF binding sites enriched (26 and 140 respectively). These TFs with enriched sites are likely important for regulating high salinity cell-type responses.

To further investigate the extent the current knowledge of large-scale TF binding data can explain high salinity up-regulation in these root cell types, we built machine learning models using the presence of CIS-BP (24) or DAP-seq (25) sites in the promoter of a gene as predictors of whether the gene in question would be up-regulated in a particular root cell-type or not (**Figure 2**). This approach allowed us to integrate information from all *in vitro* TFs into one model that could identify informative TFs that were predictive of expression patterns. Here the machine learning model performance is measured using the Area Under Curve-Receiver Operating Characteristic (AUC-ROC), which jointly considering false positive and true positive rates, where AUC-ROC = 1 indicates a perfect model and AUC-ROC = 0.5 indicates the model is no better than random guessing. Consistent with the interpretation that a subset of the TFs are likely involved in root cell-type high salinity up-regulation, models based on CIS-BP (AUC-ROC = 0.63-0.71) or DAP-seq (AUC-ROC = 0.58-0.68) were better than randomly expected for all six cell-type predictions (**Fig. 2A**). Among these predictions, models predicting END and STE response had the best performances (for an alternative measure of performance using precision-recall curves, see (**Fig. S1A, B**). For comparison, similar models were also established using putative *cis*-regulatory elements (pCREs) derived from whole organ or cell-type transcriptome (**Fig. 2B**). They will be discussed in later sections.

**Fig. 2.**
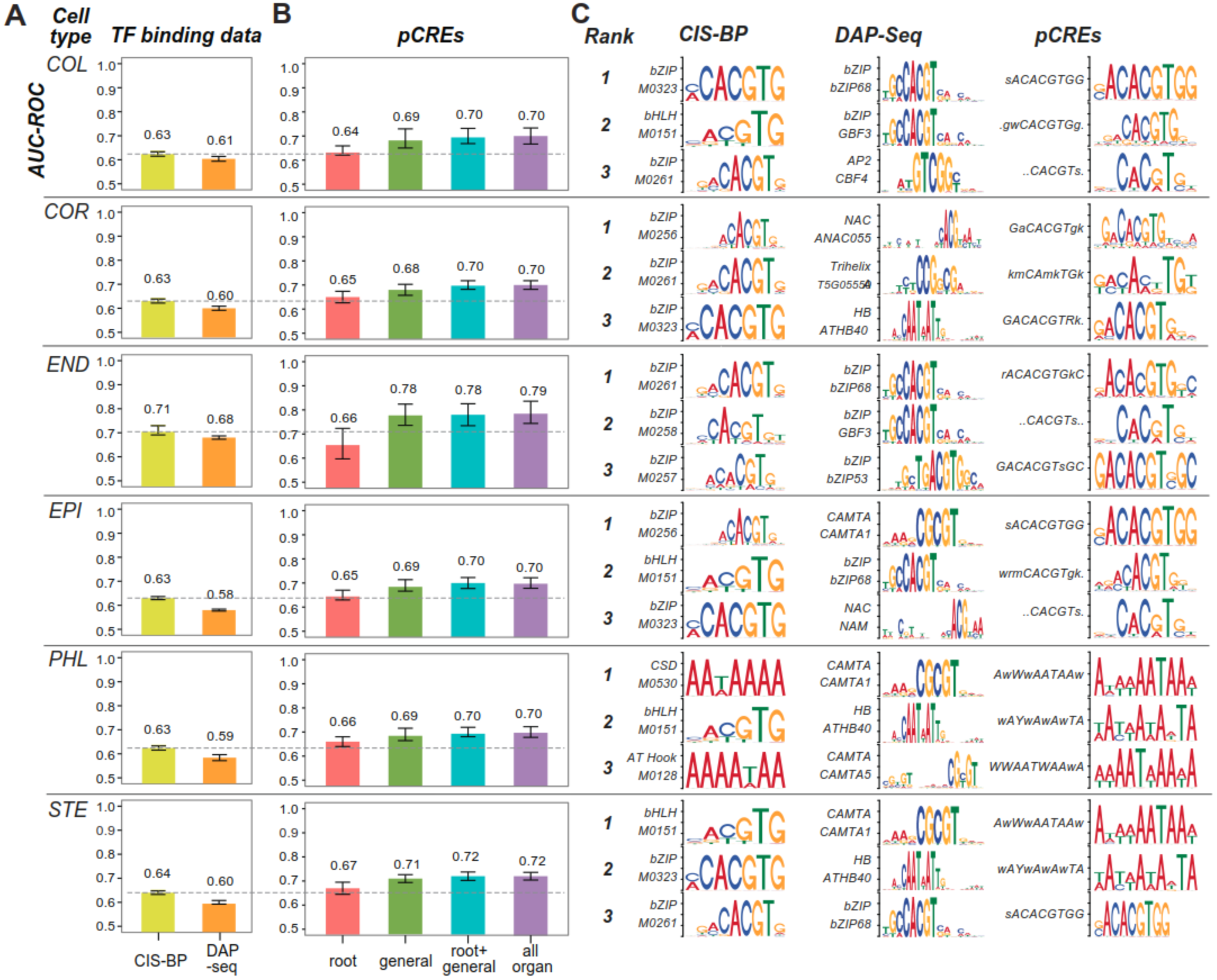
Performance of cell-type high salinity up-regulation prediction models using in vitro TF binding data and organ pCREs. **(A)** Barplot of AUC-ROC values of prediction models using CIS-BP (yellow) and DAP-seq (orange) data. **(B)** Barplot of AUC-ROC values of prediction models using organ pCREs: root (pink), general (green), union of root and general (blue), and all organ (root+general+shoot; purple) pCREs. **(C)** Top three CIS-BP and DAP-seq motifs as well as top three pCREs based on the importance score of machine learning predictions.

Next, we asked what the most important TFs were for predicting cell-type high salinity response based on the importance scores of machine learning models (see **Methods**). While some TFs most important for predicting high salinity response were important across multiple cell types, others were important for only one cell-type (**Fig. 2C**). For example, TFs belonging to the bZIP and bHLH TF families were repeatedly identified as important for multiple cell types. In contrast, CSD and AT-hook family TFs were important for predicting PHL only, suggesting their roles in cell-type specific regulation. We should emphasize that, although the root cell-type responsive gene expression can be predicted better than random by the *in*-*vitro* TF binding data, there is still substantial room for improvement (**Fig. 1A** and **Fig. S1**). One potential reason is that the binding data accounts for ~50% of known *A. thaliana* TFs, even though it is the most extensive for any plant species. Thus, some TFs and their associated binding sites important for regulating cell-type high salinity up-regulated response may be missed by this approach.

### Cell-type high salinity up-regulation prediction based on putative cis-regulatory elements identified at the organ level

To determine if accounting for TFs with no available *in*-*vitro* binding data will further improve cell-type high salinity up-regulation prediction, we used a set of organ (root and shoot) putative *Cis*-Regulatory Elements (pCREs) identified based on gene co-expression under high salinity as well as other stress treatment expression dataset (32) in the predictions. Three sets of organ pCREs were considered: 1) general organ pCREs -with sites that are enriched among both root and shoot high salinity up-regulated genes, 2) whole root pCREs -with sites enriched among genes up-regulated in root only, and 3) whole shoot pCREs -with sites enriched among genes up-regulated in shoot only (32). Considering that the high salinity-treated root cell types, and high salinity and osmotic stress-treated whole root have similar expression profiles (PCCs ≥ 95^th^ percentile PCC from all pairwise sample comparisons; **Fig. 1**), sites of whole root and general organ pCRE sets combined might explain root cell-type high salinity up-regulation.

Using the presence of sites of organ pCREs as predictors for machine learning, we found that the models based on whole root + general pCREs outperformed models using *in*-*vitro* TF binding data in predicting cell-type high salinity up-regulated genes (AUC-ROC = 0.68-0.78; cyan, **Fig. 2B;** example precision-recall curves in **Fig. S1C, D** for END and STE). Interestingly, there was little improvement from the TF-binding models using only whole root pCREs (red, **Fig. 2B**). Instead, the major contributors were the general organ pCREs, as those pCREs did significantly improve model performance when used alone. Finally, the addition of the third sets of pCREs, the whole shoot pCREs, to the whole root + general pCREs did not improve performance further.

We hypothesize that general organ pCREs were better predictors of cell-type high salinity response than whole root pCREs for two reasons. First, multiple representatives and derivatives of known stress responsive elements, such as the ABA-responsive element (ABRE: ACGTGG/T) were among the general organ pCREs because these elements are associated with TFs that regulate high salinity responses across organ types (32). Therefore, these pCREs would be useful predictors regardless of the cell-type of interest. Second, because the whole root pCREs were identified using whole root expression dataset, any useful signals from specific root cell types would likely be muted or lost, indicating the need to identify pCREs based on individual root cell-type gene expression data.

### Identifying root cell-type pCREs associated with high salinity up-regulation

In earlier studies, human cell-type specific CREs were identified for expression prediction using cell-type gene expression data and other information (48, 49). To identify root cell-type pCREs that might be involved in *A. thaliana* high salinity stress, we used existing root cell-type high salinity response data from COL, COR, END, EPI, PHL, and STE (9) and whole root abiotic stress data (37). We identified 3,095 pCREs from putative promoters of genes in “high salinity clusters” (see **Methods, File S2, S3**). These high salinity clusters were coexpression clusters with over-represented number of high salinity up-regulated genes. For each pCRE *X*, if its sites were enriched among high salinity up-regulated genes of a cell type *Y*, we refer to *X* as a cell-type pCRE for *Y*. The number of pCREs for each cell-type was correlated with the number of high salinity up-regulated genes from each cell-type (PCC = 0.95, *p* = 0.005) reflecting a potential relationship between the *cis*-regulatory complexity and the extent of high salinity up-regulation in different cell types. We classified a cell-type pCRE as a: 1) cell-type specific, 2) multi-cell-type, and 2) general cell-type pCRE based on if the pCRE was identified for only one, two to five, and all six cell types, respectively (**Fig. 3A**). We found 583 general cell-type pCREs (as opposed to the general organ pCREs discussed earlier, **Fig. 2B**). In addition, between 7 and 360 pCREs were cell-type specific (**Fig. 3A**), suggesting that their roles in regulating cell-type specific up-regulation.

**Fig. 3.**
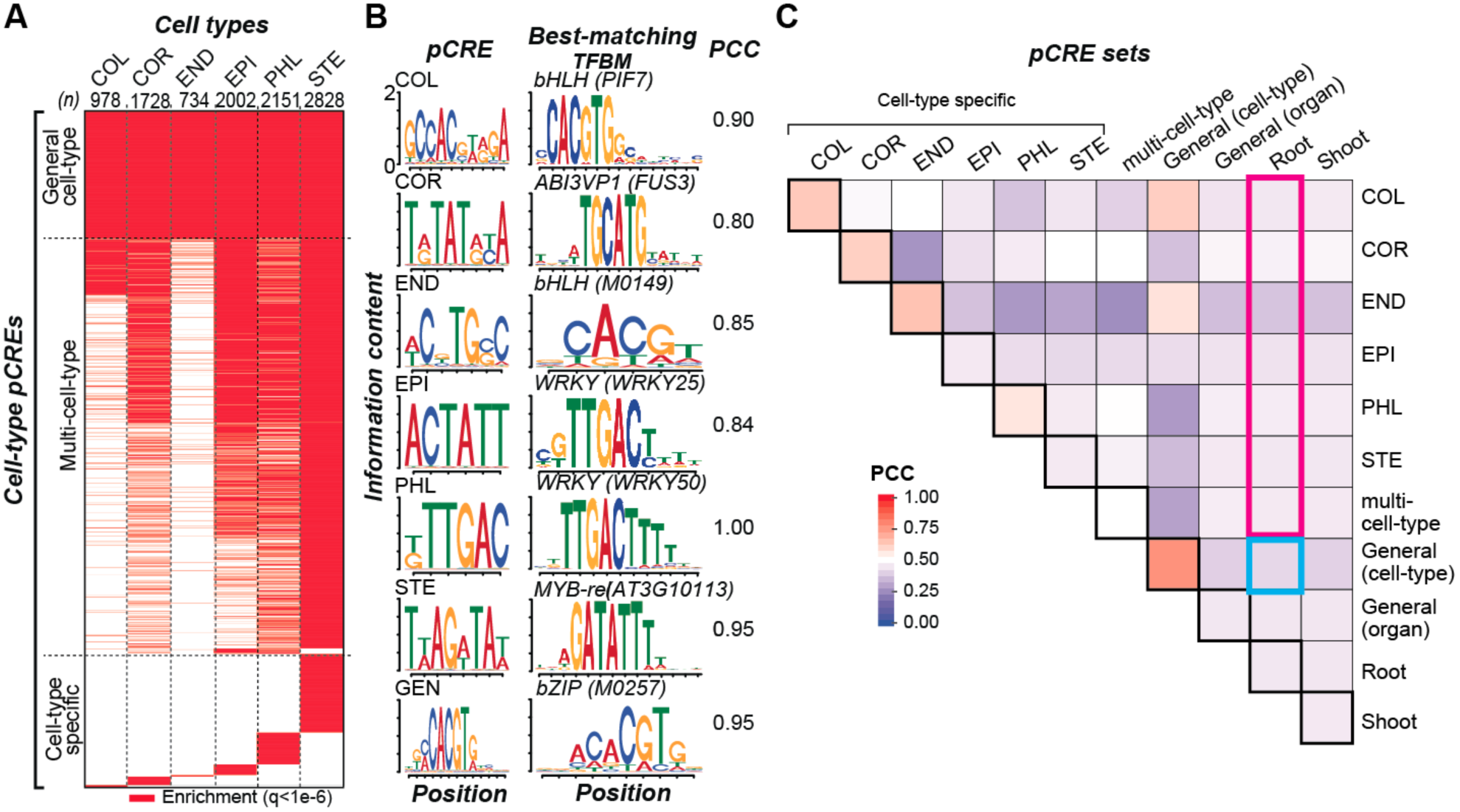
cell-type pCREs: Classification and similarity among pCRE sets. **(A)** Heatmap of over-represented pCREs. Each row is a pCRE and red color is for over-representation of that pCRE in the cell-type high salinity up-regulated genes. Top numbers: cell types and number of cell-type pCREs. **(B)** Sequence logo of the most highly enriched pCRE in each of the six cell types. For the general cell-type pCREs (GEN), an example among the highly enriched motifs is given. Fisher exact test *q*-values are COL: 1.10×10^−8^, COR: 2.94×10^−11^, END: 5.11×10^−11^, EPI: 1.24×10^−12^, PHL: 6.08×10^−14^, STE: 2.25×10^−14^, GEN: < 10^−20^) **(C)** Heatmap of similarity among pCRE sets. Similarity is calculated as PCC between PWMs. Root/shoot: pCREs enriched among up-regulated genes from the whole root/shoot data. Boxes with thicker outlines: self-self comparisons. Red and blue outlined boxes: emphasized in the text.

We identified the most significantly enriched cell-type specific pCREs for each cell-type and their best matching TF binding motifs (**Fig. 3B**). Some of these cell-type specific pCREs had high sequence similarity with a known TFBM, for example the top PHL pCRE was a perfect match (PCC = 1) to the WRKY50 TF binding motif suggesting this TF could be important in regulating PHL high salinity responsive gene expression. However, others, such as the top pCREs for COR and EPI, matched poorly to the known TFBMs and could represent novel *cis*-regulatory sequences. To assess the overall similarity between the cell-type pCREs and the organ pCREs, we determined the average similarity of pCREs (PCC) within (diagonal) and between different pCRE sets, including: 1) general organ – root + shoot, 2) whole root, 3) whole shoot, 4) general cell-type, 5) multi-cell-type (2–5), and 6) the six cell-type specific sets (**Fig. 3C**). The similarities of pCREs within four cell-type specific sets (COL, COR, END, PHL) is higher (PCC = 0.56-0.63) than the similarities across sets (average PCC = 0.41), indicating that high salinity up-regulation in these cell types involves distinct types of *cis*-regulatory sequences. Meanwhile, this is not the case for EPI and STE specific pCREs (PCC = 0.45). When we considered cross-set comparisons, one notable finding is that the general cell-type set and the whole root set (PCC = 0.42, cyan rectangle, **Fig. 3C**) are not any more similar to each other than they are to the other sets (average PCC of all cross-set comparisons = 0.44). This is also true when we compare the similarities between each cell-type specific set to the root-specific set (magenta rectangle, **Fig. 3C**). These findings further highlight the differences between whole root and root cell-type response. In addition, we are able to uncover novel *cis*-regulatory sequences using cell-type data.

### Contribution of different pCRE sets to models predicting high salinity up-regulation in different cell types

The differences between the cell-type pCREs and the organ pCREs suggest that focusing in on specific cell types will allow us to discover novel motifs important for driving high salinity up-regulation among root cell types. To assess this, for each cell-type, we first used all 3,095 cell-type pCREs as predictors to build a machine learning model (“all cell-type”) for predicting high salinity up-regulated genes in that cell-type. The AUC-ROCs for the all cell-type pCRE models ranged from 0.68 for EPI to 0.76 for END (purple, **Fig. 4A, Table S1**). Because only a subset of the 3,095 cell-type pCREs were enriched in high salinity up-regulated genes in each cell-type, we also used just the pCREs enriched in genes up-regulated under high salinity in each cell type X. The cell type X models (cyan, **Fig. 4A**) performed just as well as those using all cell-type pCREs. Although not surprising, this serves as quality control of our approach, in that adding pCREs that may be important for high salinity regulation in other cell types, does not improve model performance for that cell-type. Because the cell type X models are based on genes up-regulated in cell-type X that may also be up-regulated in one or more other cell types, such cell-type X models use a combination of three pCRE sets including: 1) general cell-type, 2) multicell-type, and (3) cell-type specific pCREs.

**Fig. 4.**
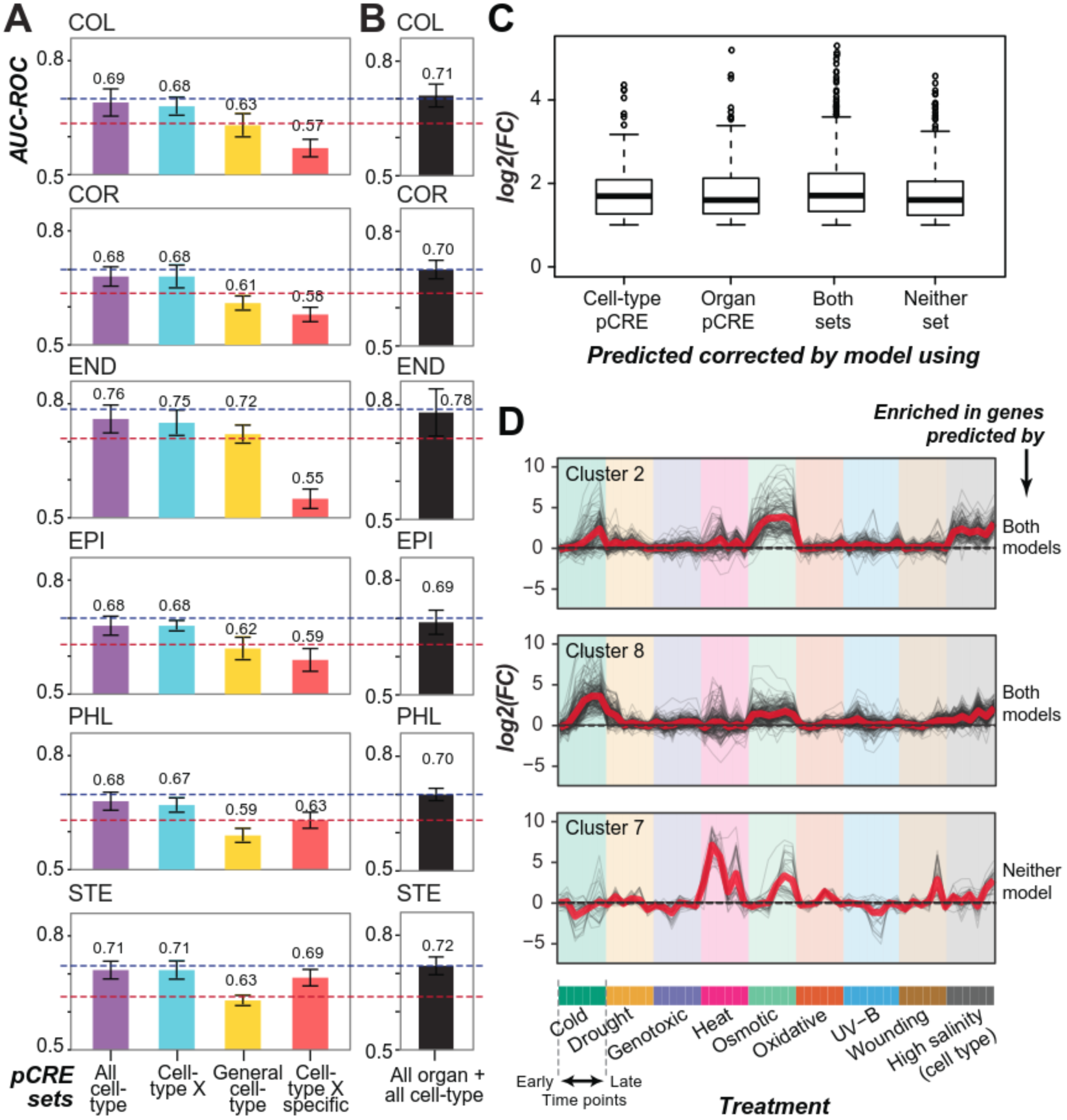
Performance of cell-type high salinity up-regulation prediction models using cell-type pCREs. **(A)** AUC-ROCs of models using four pCRE sets as predictors include: all cell-type (purple), cell-type (enriched in a cell-type X, cyan), general cell-type (orange), and cell-type specific (red). Red line: performance of the CIS-BP data-based model. Blue line: performance of the model for a cell-type using all organ pCREs. **(B)** AUC-ROCs of models for all cell types using the union of organ and cell-type pCREs. **(C)** Boxplot of log2 fold change expression values of high salinity upregulated genes in STE correctly predicted by models based on different pCRE sets. FC: fold change. **(D)** Expression profiles of STE high salinity up-regulated gene clusters (*k*-means; *k*=8) enriched in genes in the same categories as in (C). The treatment data were for whole root, except the high salinity (cell type). Gray line: individual gene. Red line: mean expression level. For each treatment, earlier time points are on the left of each color block.

To distinguish the contribution of these three sets of pCREs in the model, we next built models using only general cell-type or only cell-type specific pCREs. For COL, COR, END, and EPI, the general cell-type pCREs (yellow, **Fig. 4A, Fig. S2A**) were better predictors than the cell-type specific pCREs (red, **Fig. 4A**), indicating that in these cell types, high salinity response tends to be controlled by a general *cis*-regulatory code. On the other hand, PHL and STE high salinity up-regulation were predominantly driven by cell-type specific pCREs (**Fig. 4A**; **Fig. S2B**), highlighting the differences in general vs. cell-type specific controls between root cell types. In addition, cell-type pCREs can be used to predict cell-type high salinity up-regulated genes better than using *in vitro* TF-binding data alone (e.g. CIS-BP, red lines, **Fig. 4A; Fig. 2A**). This indicates that the cell-type pCREs further improve our knowledge of root cell-type *cis*-regulatory program.

### Characteristics of high salinity up-regulated genes corrected by models based on organ and cell-type pCREs

We next assessed if combining general organ and cell-type pCREs would further improve our ability to predict cell-type high salinity up-regulation. Surprisingly, the performance of cell-type pCRE-based models was not better than the models based on the combination of general organ and whole root pCREs in predicting up-regulation in various cell types (blue line, **Fig. 4A; Fig. 2C**). Furthermore, when we used all organ and all cell-type pCREs to build a model for each cell-type (**Fig. 4B**), our ability to predict high salinity up-regulated genes was not improved compared to using just the organ pCREs (blue line, **Fig. 4A; Fig. 2C**). This was unexpected because these cell-type pCREs were derived directly from the cell-type expression datasets and were different compared to organ pCREs (**Fig. 3D**). One explanation for the similar performances of organ and cell-type pCRE-based models was that they correctly predicted different sets of high salinity-up-regulated genes. Using the STE high salinity up-regulated genes as an example, we found that the all organ pCRE-based, and the cell-type pCRE-based models had very similar true positive rates at 61% and 63%, respectively. However, 12% of the STE high salinity up-regulated genes were only correctly predicted by the organ pCREs, and 14% were only predicted correctly by cell-type pCREs. While genes predicted correctly by both cell-type and organ pCREs had significantly higher STE expression than those not predicted correctly by either (KS test; *p* = 5e-4, **Fig. S3**), the effect size was small. In addition, there were no significant expression level differences observed in genes correctly predicted by only the cell-type or only the organ pCRE sets (KS test; *p* = 0.15-0.46) (**Fig. 4C**). However, there are differences in expression patterns and functions between these sets of genes that could shed light on why they were or were not correctly predicted (**Fig. 4D, Fig. S4**, **Table S4**).

First, STE genes correctly predicted by both the cell-type pCRE and organ pCRE-based model tend to belong to expression clusters highly up-regulated at all time points under osmotic stress and at later time points under cold stress (clusters 2 and 8, **Fig. 4D**). Notably, genes not correctly predicted by either model tend to be highly upregulated under heat stress and at later time points under osmotic stress (cluster 7, **Fig. 4D**). In addition to significant differences in expression patterns, STE genes predicted by both organ and cell-type pCRE models were enriched for genes in GO categories including response to water deprivation, ABA, and cold compared to STE genes not predicted by either pCRE sets (FET; *q* = 7~9e-3; **Table S5**). Together, this suggests that both organ and cell-type pCREs were better able to identify STE high salinity up-regulated genes that were also up-regulated at the organ level under similar conditions.

Because genes predicted by different pCRE sets were not enriched for genes with similar pCRE profiles (**Fig. S5**, **Table S6**), it remains unclear what *cis*-regulatory information lead to the difference in predictive ability. The lack of improvement in the models including both organ and celkype pCREs (**Fig. 4B**) is likely due to overfitting, where increasing the number of predictors (pCREs) does not make the model better because there is no corresponding increase in observations (high salinity up;regulated and non;responsive genes). Nonetheless, we should emphasize that, despite the caveats, the organ set and the celkype sets contain complementary *cis*-regulatory information. Jointly they provide the first glimpse of *cis*-regulatory control at the celkype level under an environmental perturbation in any plant species.

## Discussion

The existing root high salinity transcriptomic data (9) provides a rich resource to not only dissect the extent of gene expression in distinct environments across different cell types, but also provide insights into the molecular mechanisms regulating celkype transcriptional response. In this study, we identified pCREs likely responsible for celkype high salinity up;regulation in *A. thaliana.* Taking on step further, we established celkype *cis*-regulatory codes with these pCREs that reveal the relative importance of different pCREs in regulating high salinity up;regulation in different cell types. By contrasting the *cis*-regulatory codes governing whole organ (root or shoot) expression (32) with those for the cell types used in this study, we found that celkype and whole root pCREs regulate only a partially overlapping set of high salinity up;regulated genes. We also demonstrated that the pCRE;based models perform better than existing *in vitro* TF binding data in predicting high salinity up;regulation. Our findings demonstrate the feasibility of using computationally identified pCREs to establish machine learning models that can predict genome;wide transcriptional changes to specific environmental conditions at a celkype level resolution.

With the significance noted, we also become aware of a few limitations through this study. The first limitation is related to our approach in identifying pCREs from clusters of co; expressed genes. Although the approach has been fruitful, the correlation between co; expression and coagulation is far from perfect (50). In addition, not all regulatory sequences among co;regulated genes can be efficiently identified by motif finders, mainly due to the discovery that the three dimensional structure adopted by the regulatory sequences can be more important than the primary sequences (51,52). The second limitation is related to how pCREs are used for modeling. The identified pCREs are in the form of position weight matrices (PWMs) that are mapped to *A. thaliana* genome. To counter the high false positive rate in site mapping using PWMs (53), we have set a relatively stringent threshold mapping *p*-values that errs on the side of missing relevant sites. In addition, in this study the *cis*-regulatory code is built on relatively simple regulatory logic; how the presence or absence of pCRE sites in the proximal promoter region may predict up;regulation. Given the complexity of gene regulatory network, future studies incorporating hypothesized regulatory network motif information and/or considering combinatorial relations (32) between and copy numbers (54) of pCREs may further improve prediction.

The third limitation relates to the types of information considered in the model. The current model is built on celkype gene expression data with only one time point. Recently, a dataset consisting of multiple high salinity treatment time points across four root cell types (COL, COR, EPI and STE) has become available (16). It is anticipated that the time;course data will allow better clustering of co-regulated genes, and should therefore be considered in future studies. In addition to expression data, our model considers only *cis*-regulatory sequences. It is expected that incorporating additional regulatory information with a cell-type level resolution, e.g. TF binding, chromatin accessibility, and post-transcriptional regulatory data, will lead to further improvement of the regulatory model.

Apart from the limitations noted above, our study provides a comprehensive *cis*-regulatory code controlling transcription at the cell-type level in response to a stressful environment. This is a significant step forward beyond earlier predictions based on organ level transcriptional response to high salinity (32). The computational models provide estimates on how well *cis*-regulatory sequences alone may account for the regulatory information necessarily to control cell-type transcriptional response to a stressor. The models also provide mechanistic insights on how cell-type transcriptional responses are regulated by *cis*-regulatory sequences. Our study represents an important first step in establishing detailed, statistical models of stress responsive gene expression of plant cell types. With future infusion of additional regulatory information, we anticipate that the machine learning approaches used here will allow for more accurate models of spatial gene regulation in diverse environmental contexts.

## Availability

Programs and analysis pipeline relevant to this study are available on our lab GitHub repository (https://github.com/ShiuLab).

## Supplementary data

**Fig. S1. Precision/recall of END and STE high salinity up-regulation prediction models using TF binding data and organ pCREs**.

**(A)** Precision/recall curves of END high salinity up-regulation models using CIS-BP (yellow) and DAP-seq data (orange). **(B)** Same as (A) but for STE high salinity up-regulation. **(C)** Precision/recall curves of END high salinity up-regulation models using root (pink), general (green), union of root and general (blue), and all organ (root+general+shoot; purple) pCREs. **(D)** Same as (C) but for STE high salinity up-regulation.

**Fig. S2. Precision/recall of END and STE high salinity up-regulation prediction models using cell-type pCREs and union of cell-type organ pCREs**.

**(A)** Precision/recall curves of END high salinity up-regulation models using cell-type (pink), general (green), union of cell-type and general (blue), and all cell-type (purple) pCREs. **(B)** Same as (A) but for STE high salinity up-regulation. ^⋆^: depends on the predicted cell-type, ^⋆^ in (A) refers to END pCREs, in (B) refers to STE pCREs. **(C)** Precision/recall curves of END high salinity up-regulation models using **(D)** Same as (A) but for STE high salinity up-regulation.

**Fig. S3. Distribution of expression levels for STE genes predicted by both or neither pCRE sets.**

The distribution log2 fold change (x-axis) of STE high salinity up-regulated genes correctly predicted by both (red) or neither (blue) cell-type and organ pCREs.

**Fig. S4. Expression profile clusters of STE high salinity up-regulated genes.**

The expression profiles of STE high salinity up-regulated gene clusters (*k*-means; *k*=8). The treatments (X-axis) including root abiotic stress conditions as well as root cell-type specific high salinity treatment and expression levels are represented as the log2 fold change. Gray lines represent individual genes in the cluster and the red line is the mean expression level for that treatment.

**Fig. S5. Organ and cell-type pCRE profile clusters of STE high salinity up-regulated genes.**

Clusters based on the presence (black) or absence (white) of the top 100 most important **(A)** organ and **(B)** cell-type pCREs (*k*-means; *k*=8). The pCREs (X-axis) are sorted from most to least important as determined by machine learning models.

**Table S1. AUC-ROC and standard deviation values of prediction results.**

**Table S2. GO-SLIM enrichment results.**

**Table S3. DAP-seq enrichment results**

**Table S4. Enrichment of STE expression clusters in STE genes predicted by different sets of pCREs**

**Table S5. GO-SLIM enrichment results for STE genes predicted by different sets of pCREs**

**Table S6. Enrichment of STE organ cell-type pCRE profile clusters in STE genes predicted by different sets of pCREs**

**File S1. Co-expression clusters (n = 538) used for enrichment test**

**File S2. Cell-type pCREs**

**File S3. Cell-type pCRE Position Weight Matrices**

**File S4. PCC values between expression samples**

## Funding

This work was partly supported by the Fulbright Science and Technology Award to S.U.; the U.S. National Science Foundation (IOS-1546617 and DEB-1655386) and U.S. Department of Energy Great Lakes Bioenergy Research Center (BER DE-SC0018409) to S.-H.S; and NSF Graduate Research Fellowship (Fellow ID: 2015196719) to C.B.A.

## Acknowledgements

We thank Alexander Seddon for helping with expression data processing and programming in establishing the analysis pipeline during the initial phase of the project, and Ronan O’Malley for advice on the use of DAP-seq data. We also thank members of Shiu lab for their valuable suggestions to our project.

